# Priority effects determine how dispersal affects biodiversity in seasonal metacommunities

**DOI:** 10.1101/2022.02.03.479022

**Authors:** Heng-Xing Zou, Volker H. W. Rudolf

## Abstract

The arrival order of species frequently determines the outcome of their interactions. This phenomenon, called the priority effect, is ubiquitous in nature and determines local community structure, but we know surprisingly little about how it influences biodiversity across different spatial scales. Here, we use a seasonal metacommunity model to show that biodiversity patterns and the homogenizing effect of high dispersal depend on the specific mechanisms underlying priority effects. When priority effects are only driven by positive frequency dependence, dispersal-diversity relationships are sensitive to initial conditions but generally show a hump-shaped relationship: biodiversity declines when dispersal rates become high and allow the dominant competitor to exclude other species across patches. When spatiotemporal variation in phenological differences alters species’ interaction strengths (trait-dependent priority effects), local, regional, and temporal diversity are surprisingly insensitive to variation in dispersal, regardless of the initial numeric advantage. Thus, trait-dependent priority effects can strongly reduce the effect of dispersal on biodiversity, preventing the homogenization of metacommunities. Our results suggest an alternative mechanism that maintains local and regional diversity without environmental heterogeneity, highlighting that accounting for the mechanisms underlying priority effects is fundamental to understanding patterns of biodiversity.

## Introduction

Dispersal promotes the exchange of individuals between communities and thus links local population and community dynamics to regional patterns of species distributions. As a consequence, dispersal can play a key role in shaping biodiversity patterns at both local and regional scales (Kerr et al. 2002; Cadotte 2006; Grainger and Gilbert 2016). However, the relationship between dispersal and biodiversity patterns is also shaped by local conditions and processes, including biotic interactions (e.g. Kneitel and Miller 2003; Shurin et al. 2004; Altermatt et al. 2011; Carrara et al. 2012; Vanschoenwinkel et al. 2013; Catano et al. 2017). Although recent studies identify local competitive interactions as a major contributor to dispersal-diversity relationships (Pu and Jiang 2015; Lu 2021; Miller and Allesina 2021), one ubiquitous biotic process in metacommunities, the temporal sequence of community assembly, has rarely received attention.

Communities are not assembled all at once; instead, species arrive at different times and differences in the sequence of arrival times can alter the outcome of species interactions and thereby also community composition. Such priority effects (or historical contingencies) are widespread in nature and occur in systems ranging from microbes to plants and vertebrates (Alford and Wilbur 1985; Drake 1991; Kardol et al. 2013; Fukami 2015). In a metacommunity context, priority effects become important because they can alter the fate of immigrating individuals into a patch. For instance, with an early arriver advantage, individuals who arrive at a patch may either successfully establish themselves if they arrive before competitors or perform poorly and even be excluded if they arrive after competitors. By altering the outcome of local competitive interactions, priority effects have the potential to shape the relationship between dispersal and diversity patterns at local and regional scales. Although several theories predict that priority effects could affect how regional dispersal shapes biodiversity (e.g. Shurin et al. 2004; Fukami 2015; Grainger and Gilbert 2016; Miller and Allesina 2021), few have incorporated the diverse mechanisms that can generate priority effects in natural systems. Yet accounting for these mechanisms may be key to explaining why dispersal-diversity relationships are often contrary to theoretical expectations (e.g., Pu and Jiang 2015; Vannette and Fukami 2017; Toju et al. 2018).

Priority effects can be generated by at least two fundamentally different mechanisms, broadly categorized as “frequency-dependent” and “trait-dependent”. Traditionally, frequency-dependent priority effects have received the most attention and dominate the theoretical literature. They arise when fitness differences between species are small and destabilizing differences are large (i.e., positive frequency-dependent growth rates; Ke and Letten 2018). If conditions are met, the system exhibits alternative stable states, where neither species can invade when rare. Arrival time itself does not affect any per-capita rates and only matters if differences in arrival time allow for shifts in the relative frequency of species when interactions start. In contrast, trait-dependent priority effects arise when per-capita interactions between species change with their relative arrival times (Rudolf 2019). In nature, this occurs when differences in arrival time are correlated with changes in traits that affect species interactions, such as size and behavior, or modified environment from the difference in arrival times (Poulos and McCormick 2014; Rasmussen et al. 2014; Rudolf 2018; Blackford et al. 2020). In contrast to frequency-dependent priority effects, trait-dependent priority effects shift what outcomes are possible (e.g., from coexistence to competitive exclusion) by altering the parameter space (e.g., changes in interspecific competition) itself instead of favoring one of two possible states (Rudolf 2019; Zou and Rudolf 2020; Fragata et al. 2022).

These two mechanisms or priority effects are not mutually exclusive but operate at different temporal scales. Frequency-dependent priority effects require that arrival times are separated by several generations to allow populations to reproduce and increase before competitors arrive (Figure 1A; Fukami 2004; Grainger et al. 2019). Trait-dependent priority effects typically occur when arrival times are separated by less than one generation, e.g., when phenological differences between competitors lead to differences in traits such as body size (Figure 1B; Shorrocks and Bingley 1994; Rudolf 2018; Blackford et al. 2020). This distinction is especially important in a seasonal context where both can occur. Consider a community where species complete one generation per year (Figure 1C). Here, frequency-dependent priority effects can arise when one species colonizes a patch several years before others (a “first colonizer”; Figure 1C), but a later colonizing species may have earlier phenology (e.g., earlier germination, emergence from dormancy, or reproduction) and thus become “active” first within a year, allowing for trait-dependent priority effects in a year (an “early emerger”; Figure 1D). Therefore, both the different timing between and within years can determine the local outcomes of competition.

**Figure 1.**
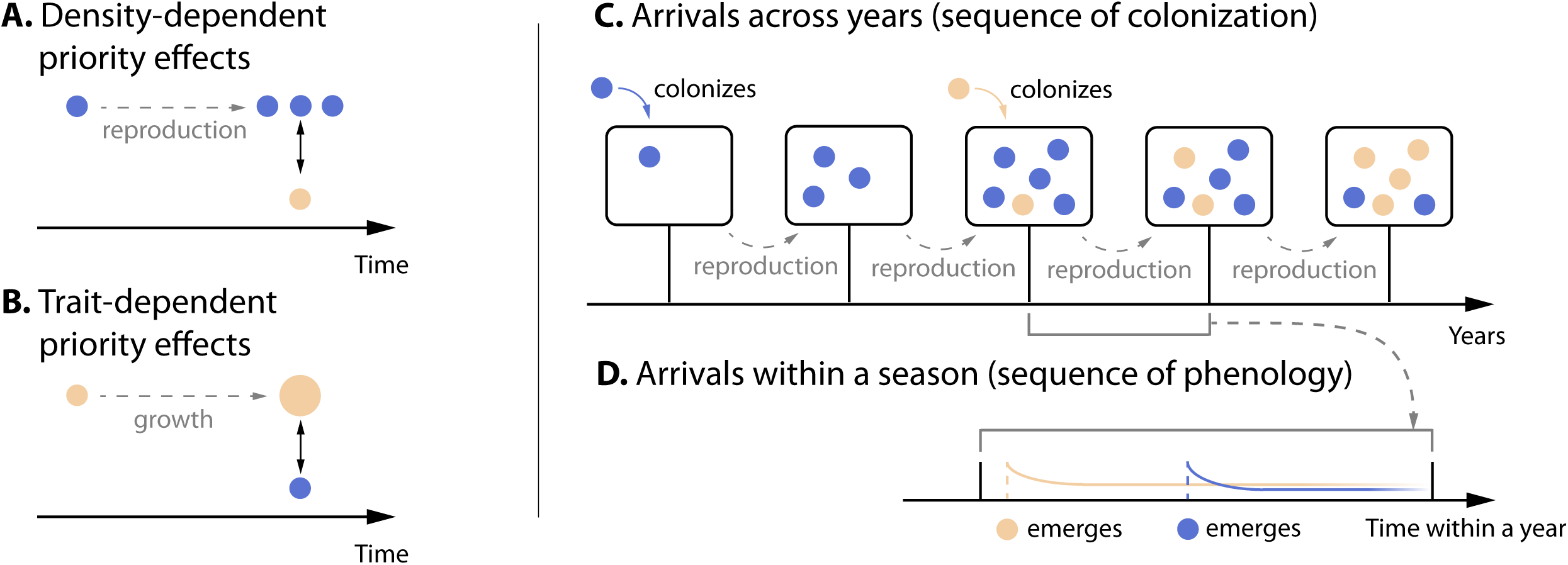
Illustration of frequency-dependent and trait-dependent priority effects and how they operate at different temporal scales. A species may arrive several seasons before its competitor, allowing it to reproduce and thus reach a higher population than the immigrating population of the late arriver (A, C). However, species can also differ in their phenology, i.e., when they become “active” (or emerge) within a season (B, D). This within-season timing can lead to changes in species interaction via changes in traits (e.g., size; B) and therefore trait-dependent priority effects. Note that a species can arrive several seasons earlier at a patch, but it can still have a later phenology within a season (C, D): the species with a frequency-dependent priority effect does not necessarily imply a trait-dependent priority effect.

These differences in priority effects in local patches have the potential to alter dispersal-diversity relationships. With frequency-dependent priority effects, positive frequency dependence could enable the more abundant first colonizer to spread over the landscape through dispersal and competitively exclude late arrivers. High dispersal will therefore decrease both local and regional diversity because one or a few regionally dominant species will eventually establish and maintain a numeric advantage in all patches, similar to the classic prediction that dispersal homogenizes (Tilman 1994; Mouquet and Loreau 2003; Leibold et al. 2004; Vanoverbeke et al. 2016; Fodelianakis et al. 2019). Following this premise, alpha (local) diversity is likely highest at intermediate dispersal while beta (dissimilarity between patches) and gamma (regional) diversity likely decreases with dispersal (Mouquet and Loreau 2002, 2003; Cadotte 2006). However, regional diversity may be retained even under high dispersal if priority effects are driven by trait-dependent mechanisms, such as when early species alter the environment or rapidly adapts to the habitat (Shurin et al. 2004; Fukami 2015; Grainger and Gilbert 2016; Leibold et al. 2019; Miller and Allesina 2021). In these cases, outcomes of competition are not only driven by relative abundances but also by within-year arrival times, or phenology. Early emergers therefore will not be excluded even if other species are more abundant. Consequences of this process should be particularly important in seasonal systems, as the frequent, periodic assembly of local communities creates many opportunities for spatial and temporal variations in arrival orders (Sheriff et al. 2011; Diez et al. 2012; Theobald et al. 2017; Carter et al. 2018; Rudolf 2019). In these seasonal systems, trait-dependent priority effects and natural variations in phenology could lead to high temporal turnover of local and regional community composition despite the homogenizing effect of dispersal.

Here, we use a spatially implicit, multispecies metacommunity model to reveal how different mechanisms of priority effects interact with dispersal to shape biodiversity at local and regional scales in competitive metacommunities. We modeled a seasonal system with spatial and temporal variations in annual emergence times (representing species phenology) and compared the observed dispersal-diversity relationships with and without trait-dependent priority effects across different initial distributions of the population. We asked the following questions: (1) How will different mechanisms of priority effects change dispersal-diversity relationships? (2) When will local and regional diversity be maintained under high dispersal?

## Methods

### Model Setup

We modeled natural habitats with seasonal disturbance, such as temperate grassland or ephemeral ponds, in which species survive periodic, unfavorable environments (e.g., winter, droughts) and reemerge at the beginning of the next growing season. We define a growing season as the growth period between a previous periodic disturbance to the next disturbance. At the beginning of the growing season (hereafter season for short), each species emerges from dormancy in a patch at an assigned time (phenology). Patches are homogeneous throughout the landscape and are not spatially structured.

Within each season, local population dynamics of species *i* in a given patch are modeled with a modified Beverton-Holt model (Rudolf 2019):

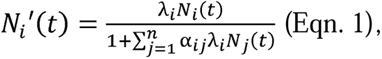

where the population at the end of time step *t* (before dispersal and end-of-season disturbance), 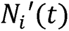, is determined by population growth of species *i*, *λ_i_N_i_*(*t*), where *λ_i_* is the intrinsic growth rate and *N_i_*(*t*) is the population of species *i* at the beginning of the season (after dispersal and disturbance of the previous season). Population growth is dependent on densities of all species with respective pairwise competition coefficients *α_ij_*. Note that this process assumes one reproduction event per season, which applies to systems such as annual plants or amphibians.

If the community is driven by trait-dependent priority effects, interspecific competition is a function of the relative emergence time:

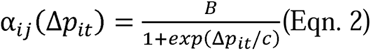

where Δ*p_it_* is the difference between actual emergence times between species *i* and *j* at the start of season *t*, calculated as *p_jt_* − *p_it_*; Δ*p_it_* > 0 when species *i* emerges first, and vice versa. *B* and *c* are constants that determine the asymptotic maximum and shape of the sigmoidal function respectively (Figure 2). When two species arrive simultaneously (Δ*p_it_* = 0), the interspecific competition coefficient is B/2. This nonlinear competition-phenology function has been validated by several empirical studies in animal and plant systems (Shorrocks and Bingley 1994; Rudolf 2018; Blackford et al. 2020). With trait-dependent priority effects, *α_ij_* is calculated based on the respective Δ*p_it_* of a given patch at season *t*. In the absence of trait-dependent priority effects, interspecific competition does not depend on the relative emergence time. We therefore calculate *α_ij_* as if Δ*p_it_* = 0: *α_ij_*(0) = *B*/2. Using Eqn. 2 in this way ensures the consistency between the presence and absence of trait-dependent priority effects. We created a competitive hierarchy across all five species by differentiating intraspecific competition (intensity of self-limitation). For each simulation, we drew intraspecific competition coefficients from a normal distribution around a mean (Table 1).

**Figure 2.**
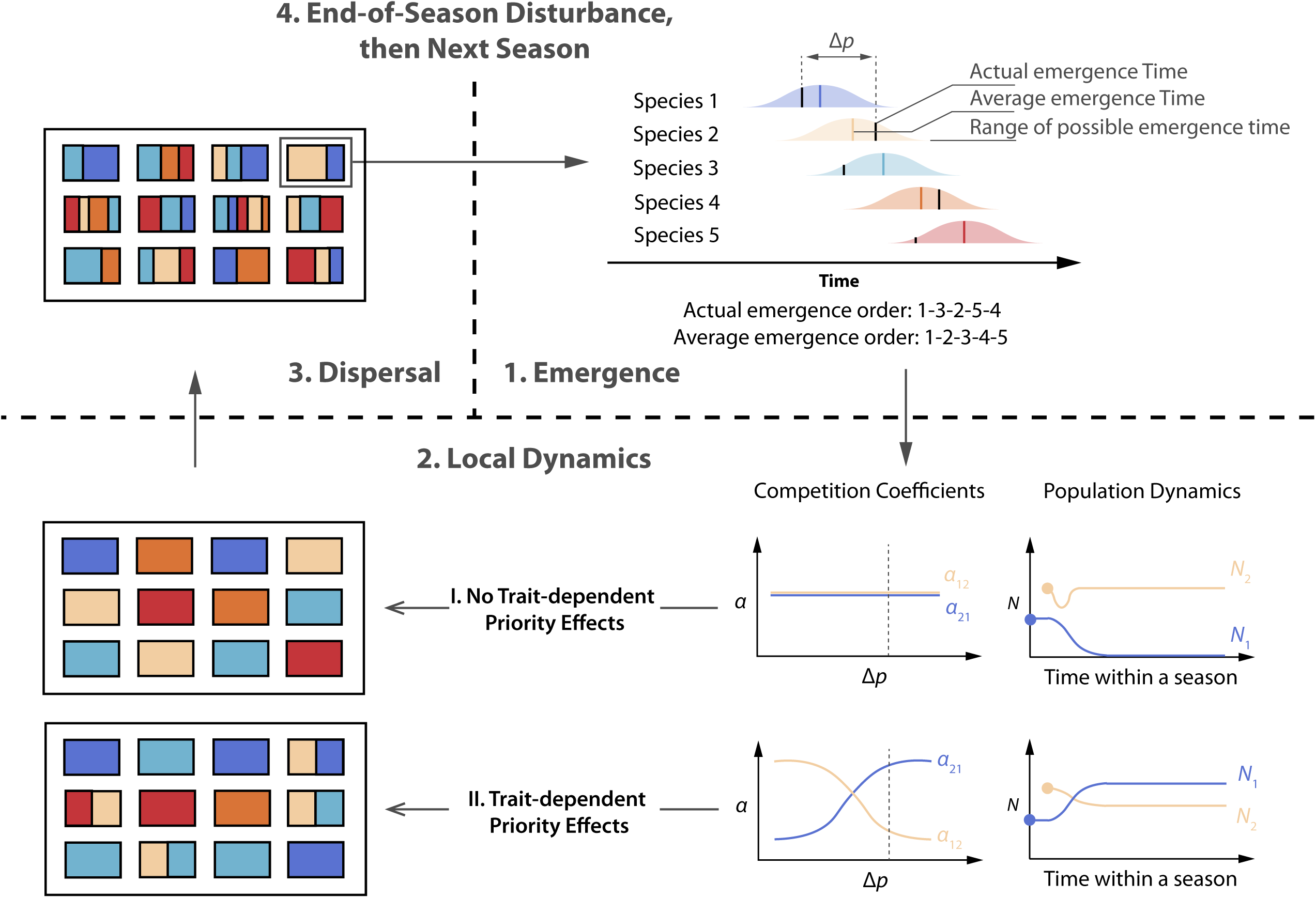
Basic processes of the metacommunity model. Colors represent species. In the hypothetical metacommunity each patch is occupied by one species and the season starts when species emerge in patches (step 1). Emergence time can vary across seasons and patches within a given species-specific range. After emergence, species compete locally (step 2) according to competition coefficients that either: (A) do not depend on the relative emergence time (without trait-dependent priority effects), or (B) depend on the pairwise difference in relative emergence time (Δ*p*) (with trait-dependent priority effects). These local patch dynamics together determine regional patterns (step 3) at the end of the season. Finally, the season ends with dispersal (step 4) and end-of-season disturbance.

**Table 1.** Major parameters and values used in simulations.

|  | Definition | Symbol | Value(s) Used |
| --- | --- | --- | --- |
| Population dynamics | Intrinsic growth rate | $\lambda$ | 100 |
| | Intraspecific competition | $\alpha_{ii}$ | Randomly drawn from normal distribution with means 0.07 (shown in main text), 0.10 or 0.15 (shown in Figure S8-S13) and standard deviation 0.01 |
| | Constants for calculating interspecific competition | $B, c$ | 0.225, -0.5 |
| | End-of-season survival | $s$ | Randomly drawn from normal distribution with mean 0.5 and standard deviation 0.05 |
| Metacommunity | Number of patches | $P$ | 50 |
| | Number of species | $n$ | 5 (shown in main text), 10 (shown in Figure S5) |
| | Number of generations in a season | $T$ | 1 |
| | Initial population | $N_0$ | 10 |
| | Dispersal rate | $r$ | Range: 0.001-0.1; representative values: 0.01 (low) and 0.1 (high) |
| Emergence time | Average emergence time | $\bar{p}_i$ | 0, 0.5, 1, 1.5, 2 for species 1-5, respectively |
| | Seasonal variation | $v$ | 2 (shown in main text), 0.5, and 8 (shown in Figure S6-S7) |

At the end of the season, species disperse in the metacommunity, and all patches are subject to an end-of-season disturbance. We modeled the dispersal process as follows: first, emigrating populations of each species in each patch are determined by the number of successes of a binomial draw with 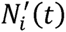 (rounded to the nearest integer) trials and probability *r* (Thompson et al. 2020), and *r* represents the dispersal rate. Emigrants are then pooled and randomly redistributed to all patches. This spatially implicit method assumes that dispersal is not affected by distance or patch choice. Spatially implicit models have been widely used in theoretical studies (e.g. Mouquet and Loreau 2002; 2003) and represent a group of empirical studies that manually transfer individuals between patches (Pu and Jiang 2015; Grainger and Gilbert 2016). We modeled the end of the season by a reduction in survival (*s_i_*), as it usually represents a period when individuals face harsh environmental conditions (drought, cold). We allowed *s_i_* to vary stochastically across seasons and patches by drawing it from a normal distribution with a mean of 0.5 and a standard deviation of 0.05. Each species then emerges at its assigned times in the next season.

### Generating Emergence Times

We randomly selected the emergence time (phenology) of species *i* from a normal distribution with mean 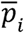 and standard deviation ν. We separated 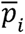 of each species by a constant interval along a gradient, with species 1 emerging on average the earliest and species 5 the latest (Figure 2). The average phenology 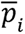 of each species is fixed, meaning that on average, the order of emergence within a season is fixed. However, the actual phenology is drawn from a normal distribution with average 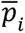 and standard deviation ν, allowing for shifts in the order of emergence. We avoided negative phenology values by normalizing the earliest phenology to 0. We repeated this process each season and for each patch and species (Figure S1). If ν > 0, this process creates variations in the phenology of co-occurring species and mimics the environmental stochasticity across space and time in natural systems (Sheriff et al. 2011; Diez et al. 2012; Theobald et al. 2017; Carter et al. 2018).

Based on field and experimental data, the phenological variation (*v*) can be up to 5-10 times the pairwise phenological differences between interacting species (Menzel et al. 2006; Kharouba et al. 2018; Rudolf 2018). Because emergence times are in a sequence with fixed intervals in our model, the pairwise differences depend on the specific species combination. To encompass possible scenarios of natural variation, we explored a wide range of *v* and picked three representative values, 0.5 (smallest pairwise difference in 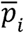), 2 (largest pairwise difference in 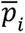), and 8 (four times the largest pairwise difference in 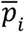), corresponding small, medium, and large variations (Table 1). We focused on results with *v* = 2.

### Initial Scenarios

To identify the different and combined effects of each type of priority effects (frequency- vs. trait-dependent) we examined the model under two different starting scenarios. The first scenario starts each patch with all five species at the same initial population (“Equal Initials”). The second scenario assumes that the first arriving species colonizes and establishes in a patch for several seasons before others arrive (“First Colonizer”). We created the latter scenario by running simulations with only one species per patch and no dispersal for 10 seasons (burn-in), with each species occupying equal numbers of patches. This ensures that the first colonizer or the “resident” species reached quasi-equilibrium abundance in the patch and only varied over time due to stochastic end-of-season survival rates before we allowed dispersal (Table 1).

We intentionally selected ranges of intra- vs. interspecific competition coefficients to allow frequency-dependent priority effects to occur in all scenarios (Table 1). However, the “First Colonizer” scenario emphasizes this frequency-dependent priority effect by allowing the initial colonizer to gain a numeric advantage over rare immigrants, while the “Equal Initial” scenario establishes a baseline that only captures the effect of seasonal variations in abundance, which is present in all cases but is not the focus of our model. We then compared the resulting population dynamics and biodiversity measures with and without trait-dependent priority effects under each scenario, resulting in a total of four different starting conditions.

### Measuring Biodiversity

We examined dispersal-diversity relationships by calculating alpha, beta, and gamma diversity of the metacommunity at the end of each simulation. We calculated alpha and gamma diversity using the Simpson’s index on local patches and the whole metacommunity respectively. Alpha diversity was averaged among all patches of a metacommunity. We quantified beta diversity (dissimilarity in community composition based on numeric abundances of species) by averaging all pairwise Bray-Curtis distances between patches. We calculate these metrics for each of the 50 simulations of a given parameter combination using respective functions in the R package *vegan* (Oksanen et al. 2013). We also calculated temporal beta diversity to quantify changes in metacommunity composition over time using the R package *adespatial* (Dray et al. 2018). Specifically, we calculated the dissimilarity (*D* metric in Legendre 2019) in species composition of a metacommunity between consecutive time steps separated by 10 seasons, excluding the first 10 seasons as burn-in. A high dissimilarity index indicates that relative abundances of species vary greatly over time.

### Model Simulations

Although our model can be extended to any number of species and patches, we simulated metacommunities with 5 and 10 competing species and 50 patches. We focus on results with 5 species here to highlight the individual populations within each patch. Each simulation was run for 110 seasons, including a 10-season burn-in period; patterns such as extinction and dominance qualitatively stabilized within 25-50 seasons. We ran 50 simulations under the same set of parameters.

All simulations were performed in R version 4.1.1 (R Core Team 2021). Code and data were uploaded to Dryad (Zou and Rudolf 2023).

## Results

Overall, we observed remarkable differences in population dynamics between scenarios with and without trait-dependent priority effects. Without trait-dependent priority effects, dispersal-diversity relationships are consistent with classic predictions but are highly dependent on initial scenarios (Equal Initials or First Colonizer). With trait-dependent priority effects, high dispersal does not lower alpha, beta, and gamma diversity in metacommunities regardless of the initial scenarios.

### Scenario 1: Equal Initials

The local population dynamics immediately highlight the difference with and without trait-dependent priority effects. Without trait-dependent priority effects, all species have an approximately equal chance of dominating a patch because the competitive hierarchy is randomly determined for each simulation. This pattern is not affected by dispersal rates, which only increases the fluctuation of populations (Figure 3). We observed that the species with the largest regional population has also the highest abundance in almost all patches, indicating both local and regional dominance that increases with dispersal rates (Figure S1). We also found a sharp decline in the temporal turnover of local communities, indicating fast competitive exclusion (Figure S2). These observations clearly show that higher dispersal promotes competitive exclusion and therefore homogenizes the metacommunity, which is consistent with the previous theory.

**Figure 3.**
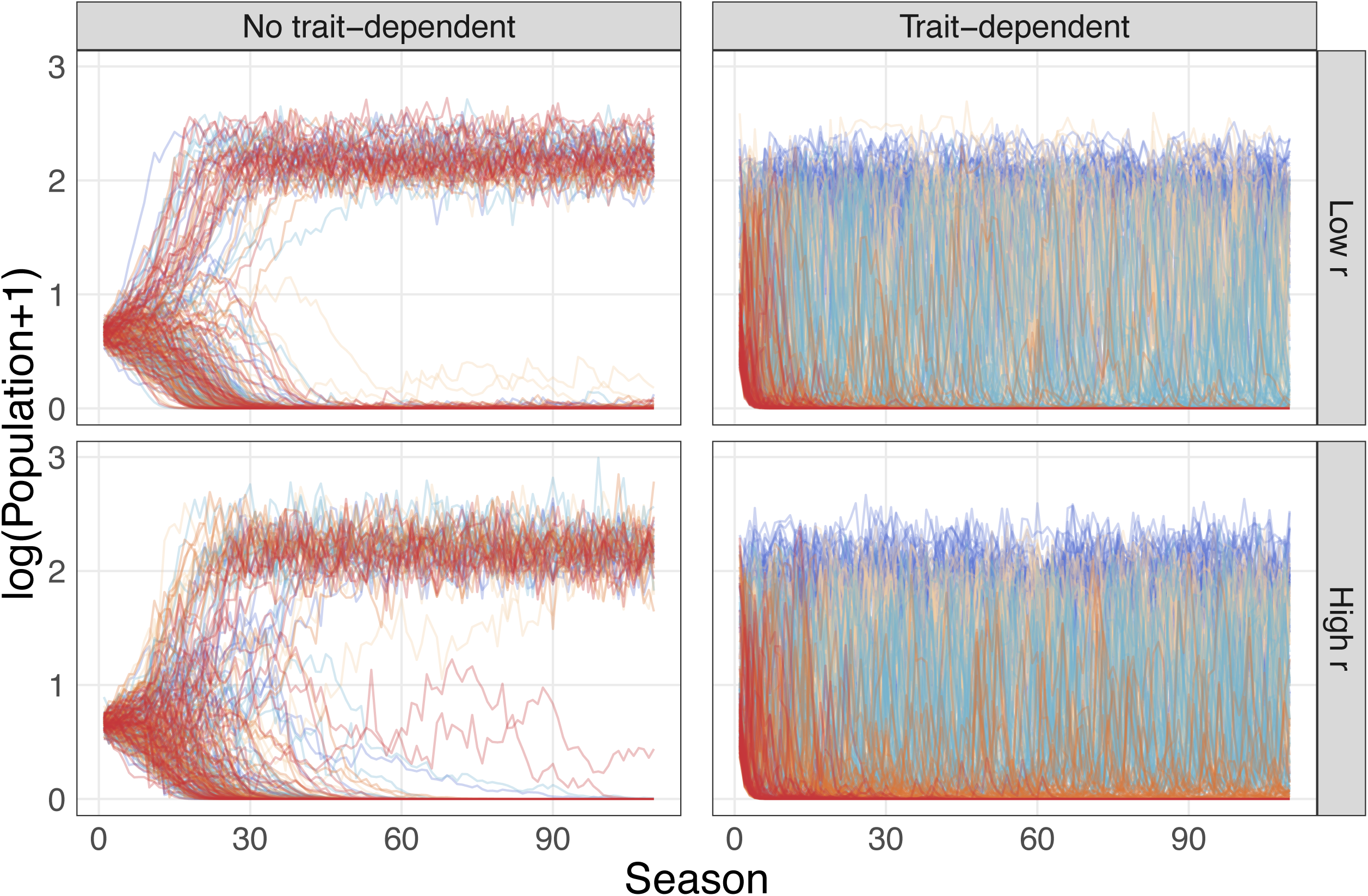
Example of population dynamics of one randomly selected patch under Scenario 1 (Equal Initials) and different dispersal rates, with and without trait-dependent priority effects. Line colors represent species. The left column shows dynamics without trait-dependent priority effects, and the right column shows dynamics with trait-dependent priority effects. Rows show results for low (0.01) and high (0.1) dispersal rates (*r*). Each line of the same color in a panel represents the population dynamics of that species from one out of 50 simulations. All population densities are transformed by natural log. See Table 1 for other parameters used.

In contrast, with trait-dependent priority effects, immigrants can successfully establish and potentially become the most abundant species in patches even at the lowest level of dispersal rates (Figure 3). This establishment is possible because the relative timing of emergence in a season determines per-capita effects and thus competitive dominance. Consequently, the species with the largest regional population does not always have the highest abundance in all patches because the rank of overall population size does not correspond to the strengths of self-limitation (Figure S1). Rather, the species that emerges the earliest also generally has the largest population (Figure S3). This also means that any species that is competitively excluded from a patch can always recolonize given sufficient seasonal variations in phenology. This recolonization is reflected in a consistently high temporal turnover in community composition within each patch (Figure S2).

Without trait-dependent priority effects, we found a weak hump-shaped pattern between alpha, beta, and gamma diversity and dispersal rates in metacommunities (Figure 5). Beta diversity is low at low dispersal rates because all local communities started with the same composition. The low alpha and gamma diversity at low dispersal rates reflects the eventual local dominance of the species with the least self-limitation. In contrast, alpha diversity increases with dispersal rates in metacommunities with trait-dependent priority effects, beta diversity remains largely constant and is not influenced by dispersal rates, and gamma diversity slightly increases with higher dispersal under trait-dependent priority effects (Figure 5). These striking differences correspond to observations of local population dynamics.

### Scenario 2: First Colonizer

Like Scenario 1, population dynamics with and without trait-dependent priority effects are remarkably different. Without trait-dependent priority effects, the first colonizer remains at high abundance in a patch at lower dispersal rates while the population of other immigrating species remains small (Figure 4; Figure S1). This pattern only changes when dispersal rates are so high that immigrating species are abundant enough to overcome the numeric advantage of the first colonizer; once this condition is met, immigrants can invade and exclude the resident (Figure 4; Figure S1). These observations are captured by temporal turnover: low dispersal rates lead to little temporal turnover of local communities that reflects the stochastic “noise” of random dispersal events. With higher dispersal rates, temporal turnover is initially high but quickly decreases to low levels because the species with the least self-limitation spreads within the metacommunity, driving others to local extinction (Figure S2). With trait-dependent priority effects, observed patterns are very similar to that of Scenario 1: the local community composition fluctuates over time (Figure S2), but the species with the earliest emergence time on average (species 1) still has the largest total population (Figure 4; Figure S4). These observations indicate that with trait-dependent priority effects, the community dynamics depend less on relative abundances but more on variations of emergence times.

**Figure 4.**
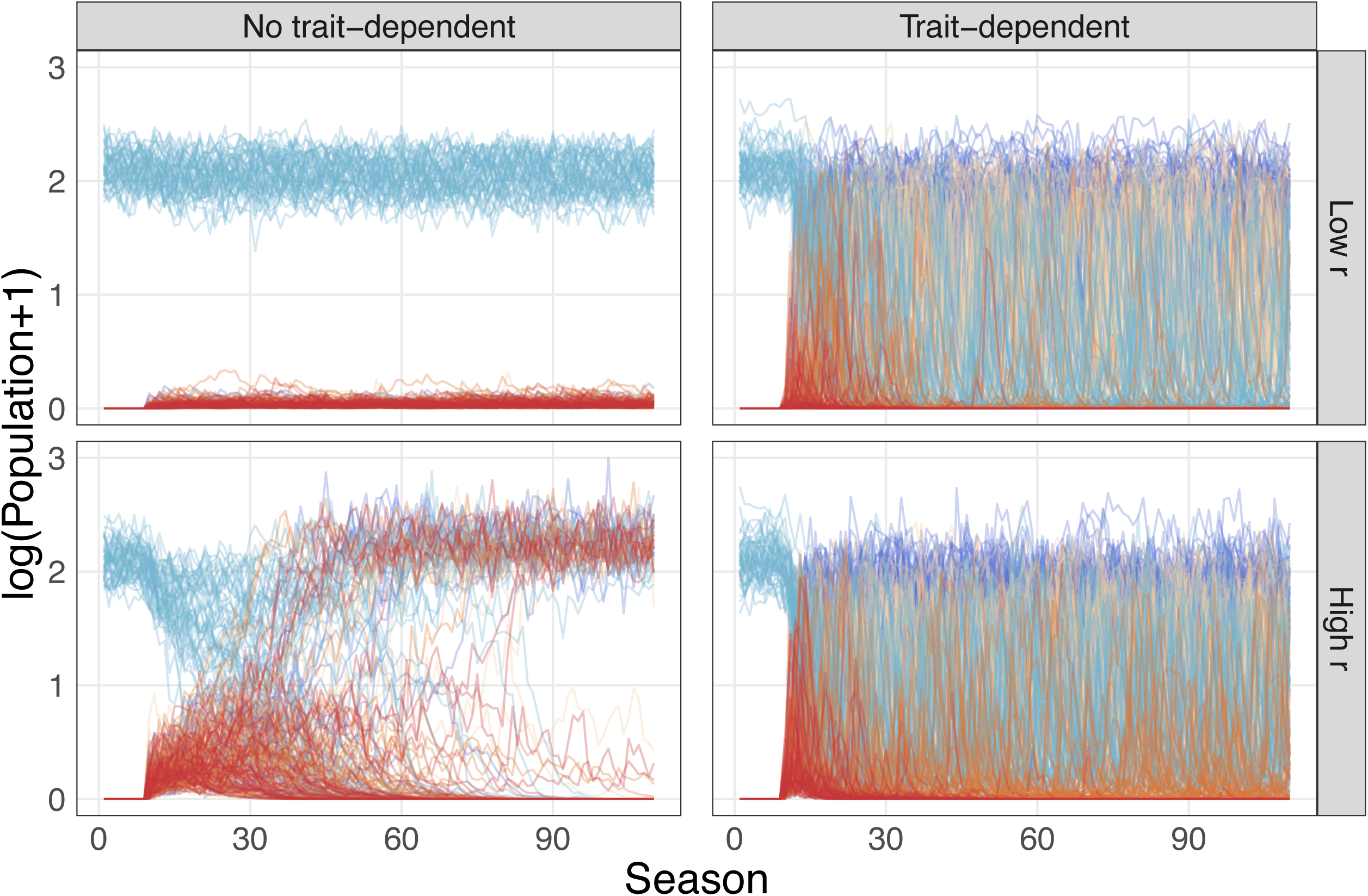
Example of population dynamics of one randomly selected patch under Scenario 1 (First Colonizer) and different dispersal rates, with and without trait-dependent priority effects. Line colors represent species. The left column shows dynamics without trait-dependent priority effects, and the right column shows dynamics with trait-dependent priority effects. Rows show results for low (0.01) and high (0.1) dispersal rates (*r*). Each line of the same color in a panel represents the population dynamics of that species from one out of 50 simulations. All population densities are transformed by natural log. See Table 1 for other parameters used.

These differences in within-patch dynamics strongly affect the local and regional dispersal-diversity relationships with and without trait-dependent priority effects (Figure 5). Without trait-dependent priority effects, alpha diversity shows a clear hump-shaped relationship with dispersal rates. Beta diversity starts highest due to both the numeric advantage and low dispersal but decreases with higher dispersal rates, indicating that dispersal homogenizes the metacommunity and leads to regional dominance of a single species. Gamma diversity decreases rapidly at higher dispersal rates as species are driven to extinction by competitive exclusion. However, with trait-dependent priority effects, all dispersal-diversity relationships are strikingly similar to those observed under Scenario 1 (compare left and right panels of Figure 5): alpha diversity increases, beta diversity slightly decreases, and gamma diversity remains the same with dispersal. Note that compared to metacommunities without trait-dependent priority effects, beta diversity is lower at low dispersal with trait-dependent priority effects because immigrants can establish in patches occupied by the first colonizer, increasing the similarity of local community composition. Overall, this shows that systems with trait-dependent priority effects are not sensitive to initial conditions, which is the opposite of systems with only frequency-dependent priority effects.

**Figure 5.**
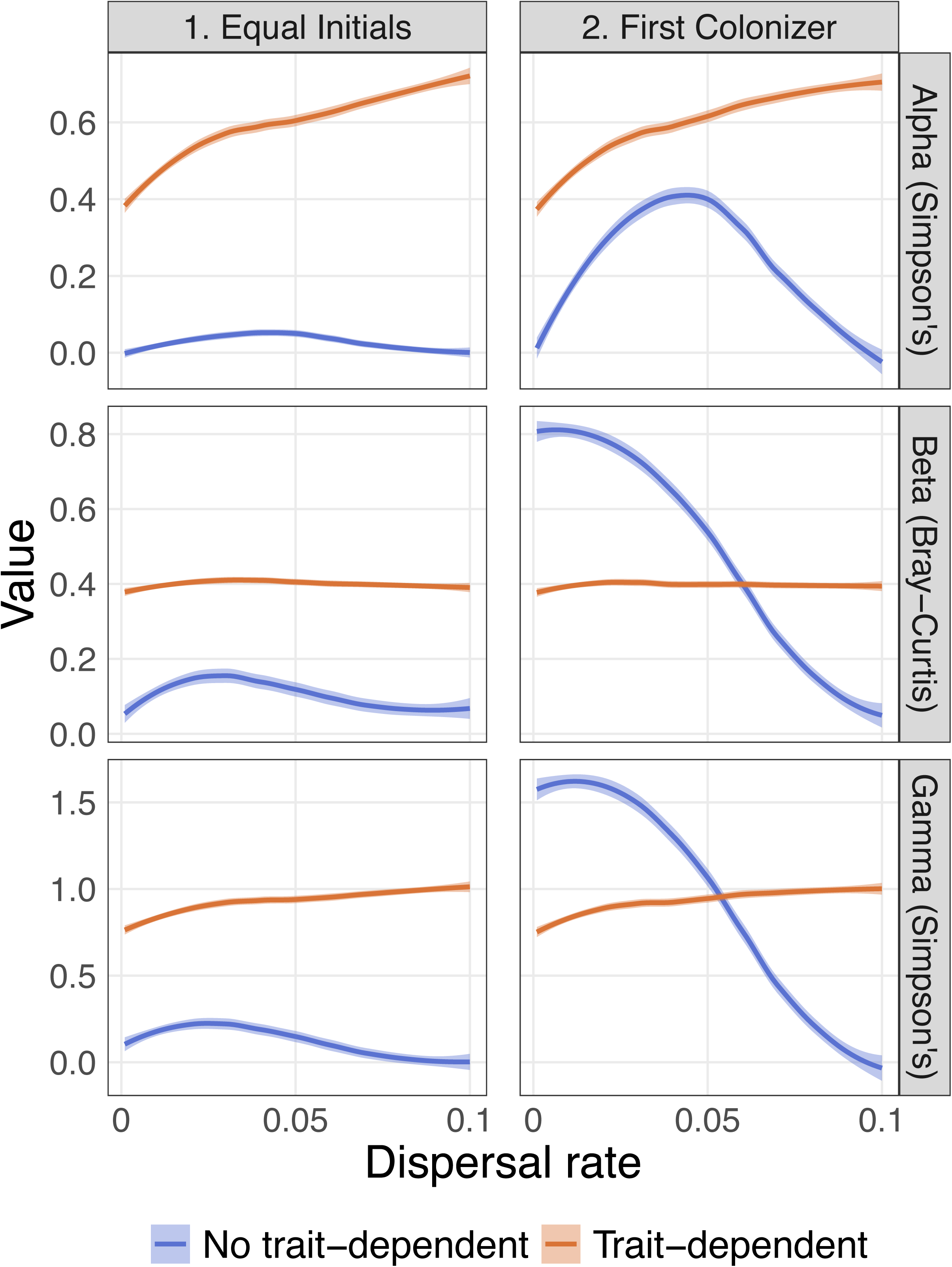
Relationship of alpha, beta, and gamma diversity and dispersal rates under the two initial scenarios, with or without trait-dependent priority effects. Left and right columns show dispersal-diversity relationships under Scenario 1 (Equal Initials) and 2 (First Colonizer), respectively. Each line is rendered from data of all 50 simulations. See Table 1 for other parameters used.

### Sensitivity Analyses

We simulated the main model with 10 species and found qualitatively the same dispersal-diversity patterns, indicating the results reflect the dynamics of more diverse systems (Figure S5). Additionally, we simulated the main model with smaller and larger variations around average phenology (i.e., the magnitude of *v*). We found that the exact dispersal-diversity relationships are strongly dependent on the magnitude of phenological variations: alpha, beta, and gamma diversity are positively correlated with phenological variation (*v*) (Figure S6, S7). This positive correlation arises because high variation increases the probability that realized phenological differences vary over time and space, thereby reducing the competitive dominance of the species which emerges 1^st^ on average. However, we still observed a consistent difference between biodiversity patterns with and without trait-dependent priority effects regardless of specific levels of phenological variations.

We then simulated the main model with five species but with two higher ranges of intraspecific competition that favor coexistence rather than frequency-dependent priority effects. We observed qualitative changes in both population dynamics and dispersal-diversity relationships without trait-dependent priority effects, but results with trait-dependent priority effects are robust (Supplementary Material, Section II). Finally, to further determine the robustness of our results we relaxed each of the following assumptions of our model: (1) each season consists of one generation, (2) dispersal happens once at the end of a season, (3) all species have equal dispersal rates (*r*), and (4) no additional spatial disturbance. Overall, we did not find a qualitative change, as we still observed remarkable differences in population dynamics and dispersal-diversity relationships between mechanisms of priority effects (results not shown).

## Discussion

Priority effects are ubiquitous in natural systems, yet their role in shaping biodiversity patterns is still poorly understood (Fukami 2015). Here we show that priority effects can play a key role in shaping the local and regional biodiversity of seasonal communities and how they are influenced by dispersal. However, the shape of this dispersal-diversity relationship depends on what type of priority effects are present. Metacommunities with only frequency-dependent priority effects are highly sensitive to the initial conditions and exhibit high spatial diversity at low dispersal rates but become homogenized as dispersal increases. In contrast, trait-dependent priority effects promote local and regional species coexistence even under very high levels of dispersal and the patterns are highly robust to variation in initial conditions. Together, these results provide novel insights into how priority effects shape the dispersal-diversity relationships in nature and highlight the importance of considering the seasonal nature of local community assembly to predict regional patterns.

### Positioning Priority Effects in Current Dispersal-Diversity Framework

Classic models predict that the dispersal-diversity relationship should be maximized at intermediate dispersal, while between-patch and regional diversity should constantly decrease with dispersal (Mouquet and Loreau 2002; 2003). Diversity peaks at intermediate dispersal because species are present in both source and sink habitats, and the population in the latter can be rescued by immigration from the former (Leibold et al. 2004). Traditionally, source-sink dynamics are expected to be driven by spatial heterogeneity in habitat conditions (Pulliam 1988). However, we still observed source-sink dynamics even in the absence of any environmental heterogeneity. In our model, the species with the highest population in a patch poses a barrier for rare immigrants to establish themselves. These patches serve as sources for the resident and sinks for all other (immigrating) species. At low to moderate dispersal rates, frequency-dependent priority effects thus maintain high beta diversity within metacommunities. At high dispersal rates, species with the least self-limitation can reach patches in large numbers and overcome frequency-dependent priority effects. The source-sink dynamics dissolve, and the homogenization effect of dispersal decreases local and regional diversity (Mouquet and Loreau 2003; Grainger and Gilbert 2016; Catano et al. 2017). Our results agree with several modeling studies involving patch preemption, which prevents immigrants from colonizing patches occupied by previous or current residents. Here, patch preemption can be considered a source of environmental heterogeneity that can arise from trait-dependent priority effects via habitat modification. Past models show that this mechanism promotes regional coexistence (Yu and Wilson 2001; Calcagno et al. 2006; Miller and Allesina 2021).

Furthermore, we found that the shape of dispersal-diversity relationship is highly sensitive to initial conditions (i.e., the presence or absence of initial numeric advantage) but only in metacommunities without trait-dependent priority effects. Notably, our scenario where all species start at the same densities (Scenario 1) coincides with a design often used in lab experiments (Grainger and Gilbert 2016), but the resulting population dynamics and dispersal-diversity relationships are different from those obtained under a scenario where a species was allowed to establish in a patch before others could colonize (Scenario 2). This result is not previously emphasized in theory (but see Lu 2021) and could at least partly help explain why the diversity-stability relationship differs among studies that use different initial settings (Grainger and Gilbert 2016). We did not observe this sensitivity in models with trait-dependent priority effects because the high spatiotemporal variation in interspecific competition determined population dynamics, masking any effect of initial conditions.

On the contrary, diversity patterns in metacommunities with trait-dependent priority effects are much less affected by dispersal or initial conditions. In these metacommunities, immigrants may successfully colonize a patch regardless of their numeric disadvantage as their colonization success is more dependent on the order of emergence. If an immigrating species emerges early enough within a season, this advantage (relative increase in interspecific competitive ability) can overcome its numeric disadvantage toward the resident species, allowing it to establish in the patch. Given the temporal and spatial variation in within-season emergence times (phenology), trait-dependent priority effects can promote local and regional coexistence of species by preventing competitive exclusion and homogenization at a high dispersal rate. Indeed, smaller phenological variations lead to a more definitive competitive hierarchy, lowering local and regional biodiversity. Our results suggest that seasonality and phenological variations are important factors promoting coexistence in spatial communities, in addition to other classic mechanisms such as spatial heterogeneity (Leibold et al. 2004; Schreiber and Killingback 2013).

The trait-dependent priority effect in our model appears conceptually similar to processes implemented by some previous models (e.g. biotic heterogeneity in Shurin et al. 2004; monopolization effects by local adaptation in Vanoverbeke et al. 2016). Yet, they found that positive feedback from early arrivers (in our model, first colonizers) enables their regional dominance. This result represents what we found in metacommunities with high dispersal and initial numeric advantage, but not in those with trait-dependent priority effects. This discrepancy arises because we incorporated seasonality and realistic spatial and temporal phenological variations in our model. These variations lead to a constant reshuffling of competitive hierarchies across time and space that reduces or prevents local and regional extinction, as predicted by previous theoretical results in non-spatial models (Rudolf 2019; Zou and Rudolf 2020). In our model, phenological variations changed interspecific competition coefficients and therefore the relative strengths between them and each species’ self-limitation. When aggregated on a spatial or temporal scale, this reshuffling of competitive hierarchies can also lead to intransitive competition, which is predicted to stabilize multispecies systems (Allesina and Levine 2011; Levine et al. 2017).

Our results provide a theoretical basis for interpreting many empirical results, especially those explicitly considering the role of priority effects. For instance, experiments in freshwater protists found that community composition in homogeneous patches was strongly correlated to the temporal sequence of assembly despite dispersal (Pu and Jiang 2015). Our results suggest that this counterintuitive pattern can be easily explained by trait-dependent priority effects. Consistent with this prediction, Pu and Jiang (2015) reported changes in interspecific competition based on arrival orders of species, indicating the presence of trait-dependent priority effects. Our observations also corroborate the result that priority effects in local nectar microbial communities help maintain beta diversity (Vannette and Fukami 2017), although we do not observe a consistent increase of beta diversity with higher dispersal, as in the experiment. Toju et al. (2018) also found a persistent numeric advantage of initially abundant species after prolonged dispersal in nectar microbial communities. This may indicate that dispersal in this empirical study was not high enough to homogenize all local patches or priority effects initiated by numeric advantage were maintained in other trait-dependent mechanisms, such as habitat modification.

### Future Directions

Phenology and priority effects are key factors that structure population dynamics and coexistence patterns in natural communities. Although the importance of priority effects is increasingly recognized (Fukami 2015; Rudolf 2019), they are rarely studied in an explicitly spatial context. Our model indicates the need to consider specific mechanisms of priority effects when studying spatial dynamics of metacommunities and vice versa. For instance, systems that display robust diversity measurements regardless of dispersal might be more influenced by trait-dependent processes. In these systems, considering specific trait-dependent mechanisms of priority effects may help to understand otherwise unexpected patterns (e.g. Pu and Jiang 2015; Vannette and Fukami 2017; Toju et al. 2018). Conversely, metacommunities with species known to display trait-dependent priority effects (e.g. amphibians, Rudolf 2018; dragonflies, Rasmussen et al. 2014; plants, Kardol et al. 2013; Blackford et al. 2020) may show different dynamics at a regional scale compared to metacommunities with frequency-dependent priority effects. Yet, few studies have considered this aspect when studying dispersal-diversity relationships in these systems, although many metacommunities are driven by stochastic processes that promote priority effects (e.g. Chase 2007).

Our model predicts that seasonal metacommunities under spatial and temporal variations in phenology may behave differently based on the mechanism of priority effects. This is relevant to many systems in nature, including ephemeral ponds, nectar microbiome, pitcher plant inquilines, rocky intertidal pools, and any other periodic communities where such variations could occur (Berlow and Navarrete 1997; Kneitel and Miller 2003; Chase 2007; Sheriff et al. 2011; Theobald et al. 2017; Vannette and Fukami 2017). Although recent studies of these systems indicate that phenology can vary considerably between communities (Sheriff et al. 2011; Diez et al. 2012; Theobald et al. 2017; Carter et al. 2018), few have addressed the local and regional level consequences of these variations. As climate change could exacerbate phenological shifts and variations (Parmesan 2006; Diez et al. 2012; Wolkovich et al. 2014; Pearse et al. 2017; but see Stemkovski et al. 2022), the arrival order of species may differ greatly over space and time, leading to increasing opportunities for priority effects. Understanding how metacommunities respond to this increasing reshuffling of arrival order (and likely competitive hierarchy) in the long term could help evaluate the responses of biodiversity to climate change.

We have extensively investigated different forms of dispersal and incorporated variations in phenology, but the influence of another aspect of spatial communities, environmental heterogeneity, remains unexplored. Heterogeneous patches could serve as a spatial refuge for some species, promoting regional coexistence (Shurin et al. 2004; Schreiber and Killingback 2013). Environmental heterogeneity may also induce trait-dependent priority effects, leading to monopolization effects (Urban and De Meester 2009; De Meester et al. 2016) or legacy effects (Miller and Allesina 2021) in local patches. Furthermore, environmental heterogeneity could covary with other factors that are coupled with another process, such as spatial variation of arrival time (e.g., species may arrive early in warmer patches), leading to more complex dynamics. Finally, we focused on systems with one generation per growing season (e.g., annual plants, amphibians), but the life history could also affect the importance of trait-dependent priority effects: with multiple generations per season (e.g., aphids, protists), frequency-dependent priority effects could arise from within-season population dynamics, leading to alternative patterns of biodiversity (Zou and Rudolf 2020). Collectively, our results emphasize that the temporal sequence of community assembly is a critical process shaping biodiversity in spatial communities. A mechanistic understanding of this process is fundamental to our knowledge of how biodiversity responds to critical environmental processes in a changing world.

## Acknowledgement

We thank members of Miller and Rudolf labs for their feedback on the project and Lydia Beaudrot, Joshua Fowler, Tom E. X. Miller, and Chuliang Song for their comments on the manuscript. Feedback from Erol Akçay, Karen Abbott and four anonymous reviewers greatly improved the manuscript. Funding was provided by NSF DEB-1655626.

## Statement of Authorship

H.-X.Z. conceived the idea, designed, and performed the research with assistance from V.H.W.R. H.-X.Z. wrote the first draft, and all authors contributed to revisions.

## Data and Code Accessibility

Data and code are deposited at Dryad (Zou and Rudolf 2023).

