## Supplementary material for "Priority effects determine how dispersal affects biodiversity in seasonal metacommunities"

### Supplementary Material for Priority effects determine how dispersal affect biodiversity in seasonal metacommunities

Accepted, *The American Naturalist*

#### Section I: Supplemental Figures

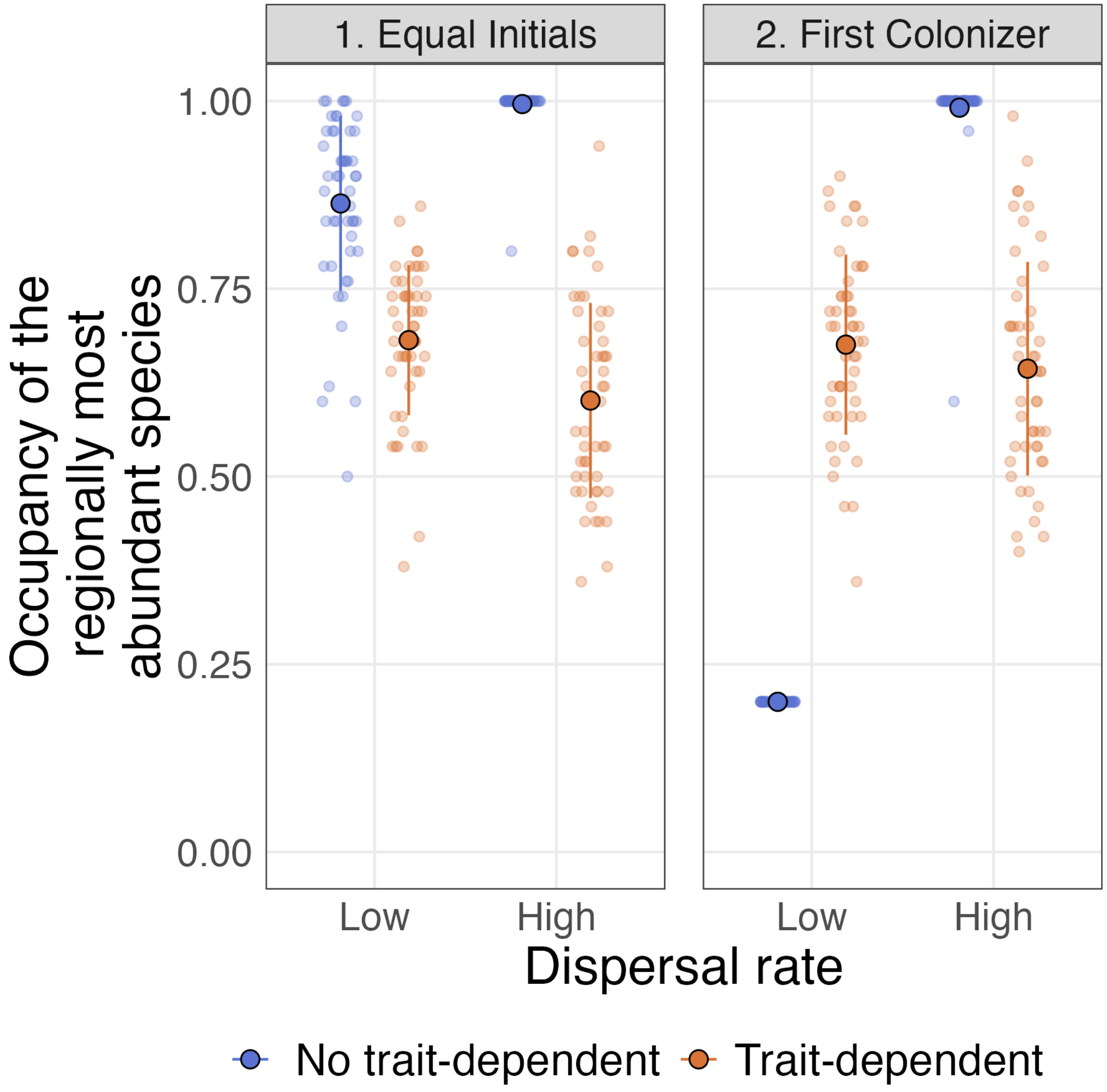


Figure S1. Patch occupancy of species with the least self-limitation, calculated as the proportion of all patches where it species has the highest population, under low (0.01) and high (0.1) dispersal rates. Left and right panels show results under Scenario 1 (Equal Initials) and 2 (First Colonizer), respectively. Solid points and error bars show the mean and standard deviation, and translucent points show actual occupancy data. See Table 1 for other parameters used.

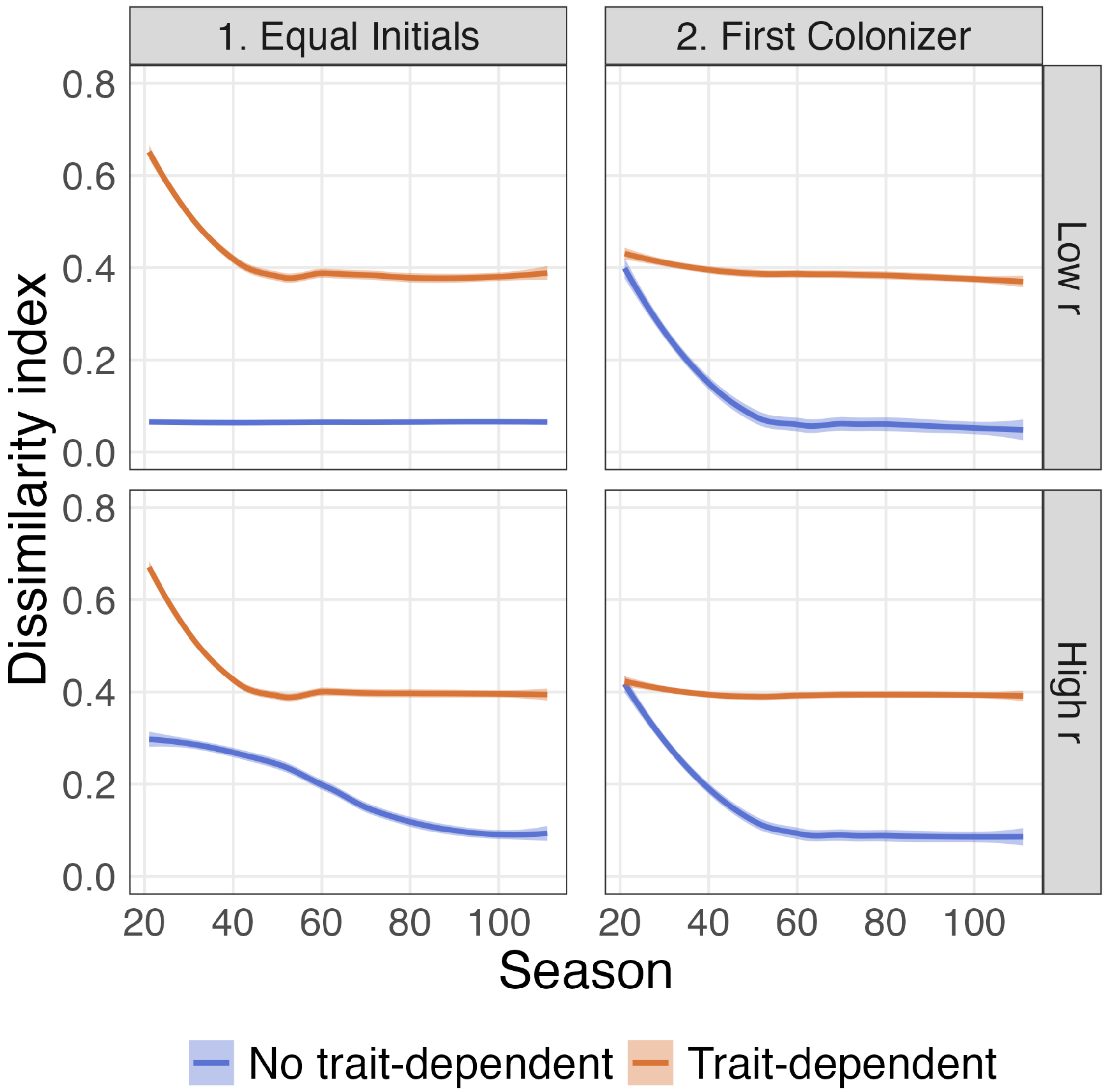


Figure S2. Temporal dissimilarity index calculated from one randomly selected simulation under Scenario 1 under different dispersal rates, with and without trait-dependent priority effects. Left and right columns show results under Scenario 1 (Equal Initials) and 2 (First Colonizer), respectively. Rows show results for low (0.01) and high (0.1) dispersal rates (r). The dissimilarity index was calculated from metacommunities between consecutive time steps separated by 10 seasons, omitting the first 10 seasons as burn-in. Each line is rendered from data of all 50 simulations. See Table 1 for other parameters used.

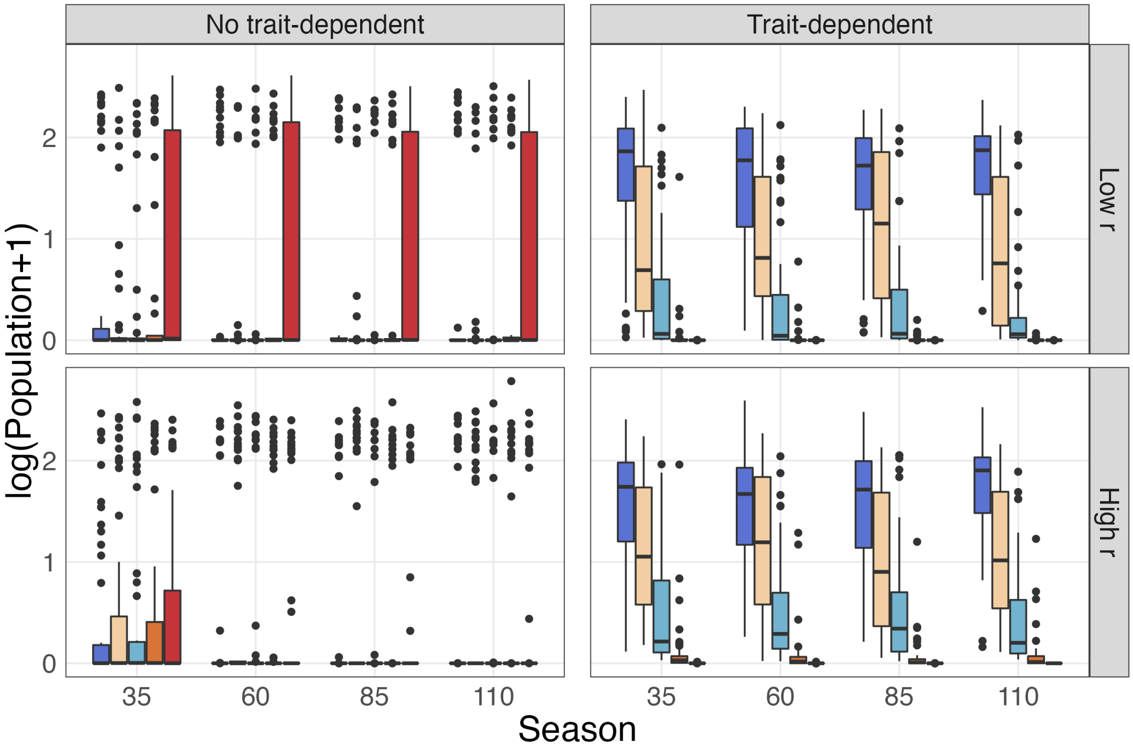


Figure S3. The regional population of each species under Scenario 2 (First Colonizer) at 35, 60, 85, and 110 seasons, low and high dispersal rates, with and without trait-dependent priority effects. The duration of the simulation is 100 seasons with a 10-season burn-in period. The left column shows dynamics without trait-dependent priority effects, and the right column shows dynamics with trait-dependent priority effects. Rows show results for low (0.01) and high (0.1) dispersal rates ($r$). Boxplots show data from all 50 simulations. All population densities are transformed by natural log. See Table 1 for other parameters used.

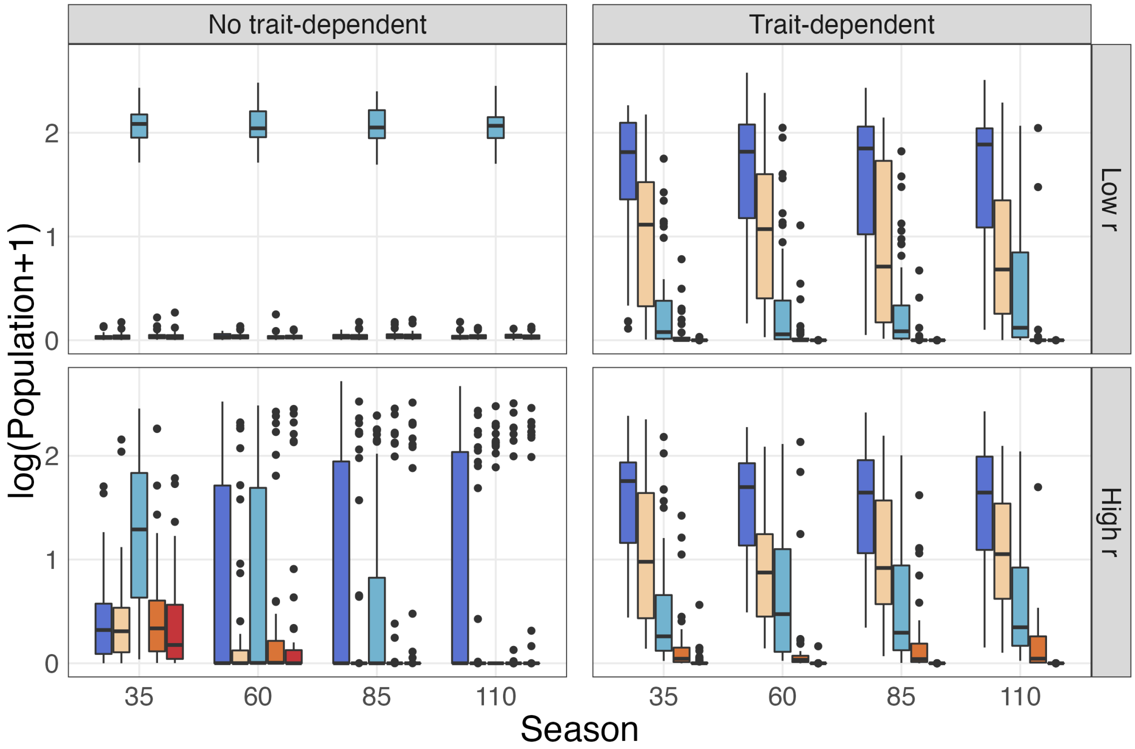


Figure S4. The regional population of each species under Scenario 2 (First Colonizer) at 35, 60, 85, and 110 seasons, low and high dispersal rates, with and without trait-dependent priority effects. The duration of the simulation is 100 seasons with a 10-season burn-in period. The left column shows dynamics without trait-dependent priority effects, and the right column shows dynamics with trait-dependent priority effects. Rows show results for low (0.01) and high (0.1) dispersal rates ($r$). Boxplots show data from all 50 simulations. All population densities are transformed by natural log. See Table 1 for other parameters used.

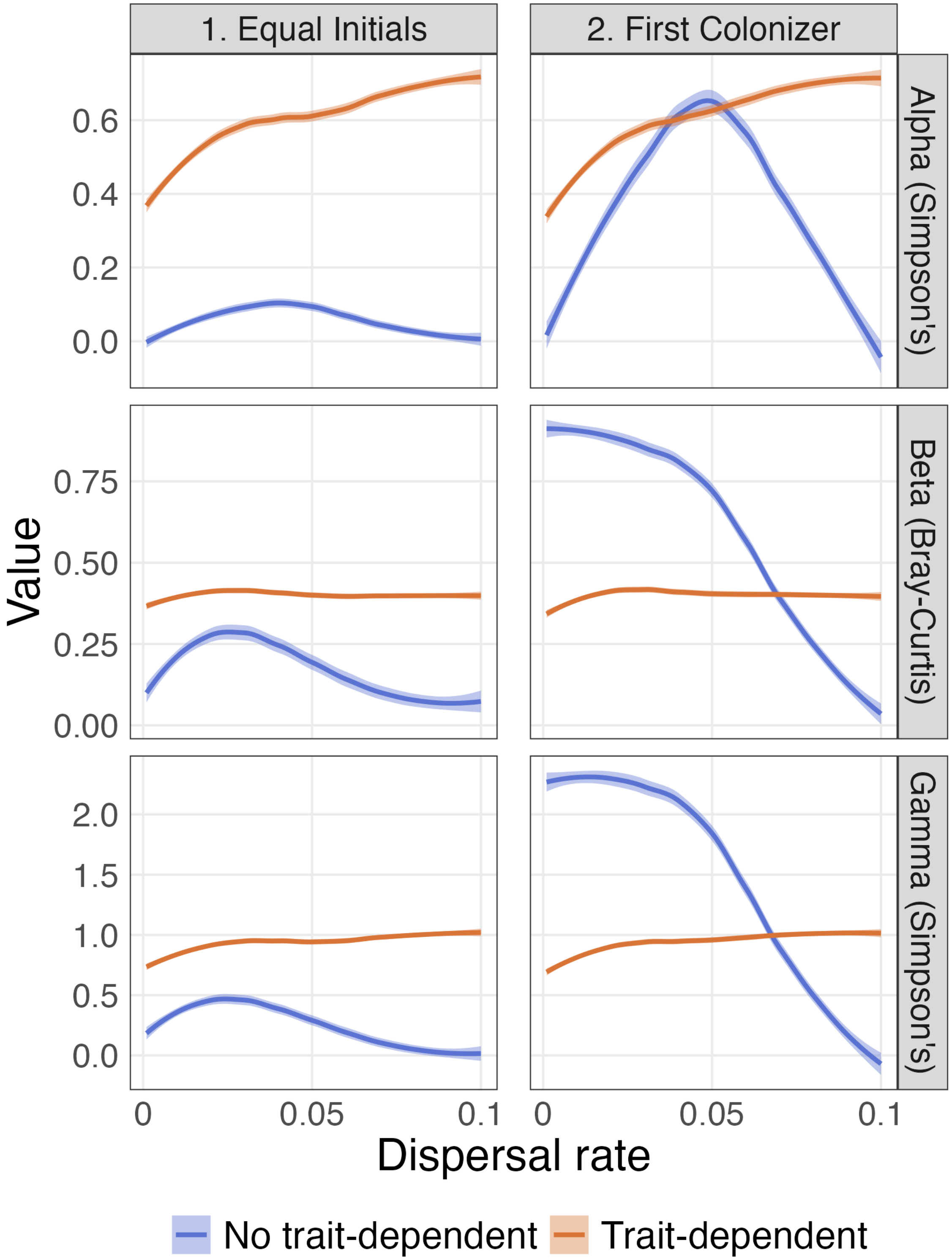


Figure S5. Relationship of alpha, beta, and gamma diversity and dispersal rates under the two initial scenarios, with or without trait-dependent priority effects, using 10 species. Left and right columns show dispersal-diversity relationships under Scenario 1 (Equal Initials) and 2 (First Colonizer), respectively. Each line is rendered from data of all 50 simulations. See Table 1 for other parameters used.

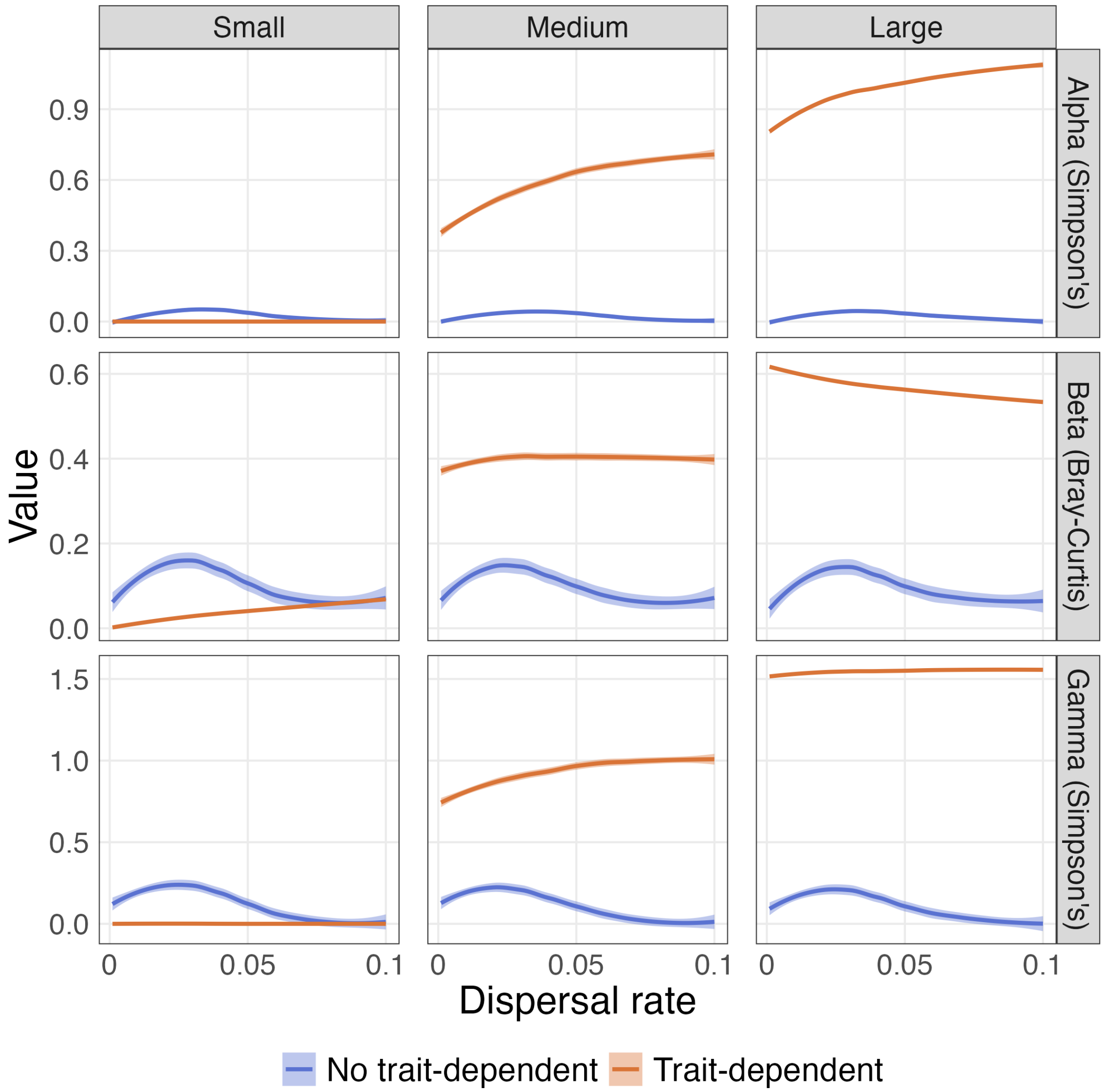


Figure S6. Relationship of alpha, beta, and gamma diversity and dispersal rates under different variations in emergence times ($v$), with or without trait-dependent priority effects. All simulations are under Scenario 1 (Equal Initials). Left, middle and right columns show results with a small (0.5), medium (2), and large (8) variation in emergence times, respectively. Each line is rendered from data of all 50 simulations. See Table 1 for other parameters used.

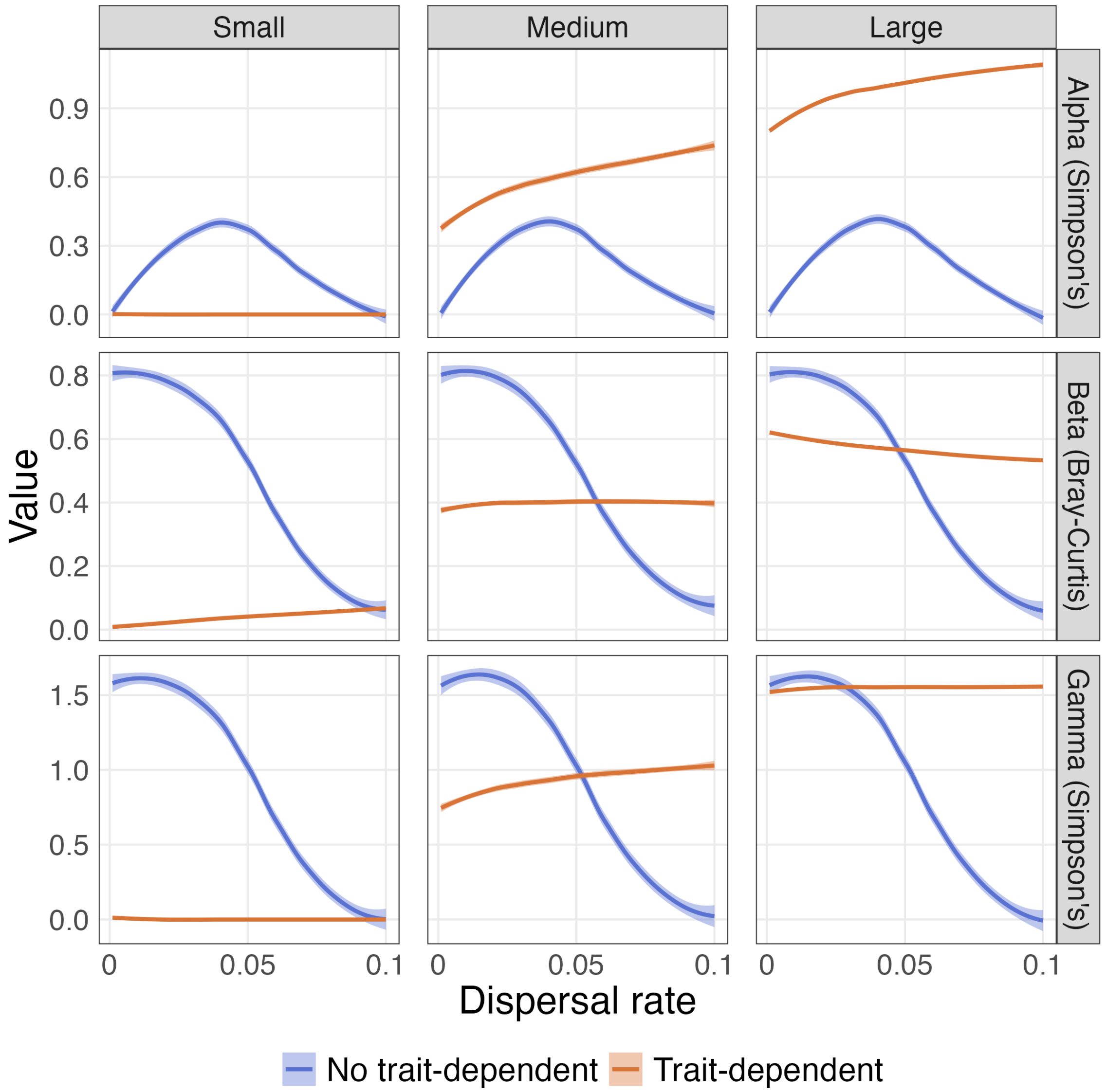


Figure S7. Relationship of alpha, beta, and gamma diversity and dispersal rates under different variations in emergence times ($v$), with or without trait-dependent priority effects. All simulations are under Scenario 2 (First Colonizer). Left, middle and right columns show results with a small (0.5), medium (2), and large (8) variation in emergence times, respectively. Each line is rendered from data of all 50 simulations. See Table 1 for other parameters used.

### Section II: Additional Simulations

To test whether differences between metacommunities with and without trait-dependent priority effects are robust to selection of intraspecific competition coefficients, we ran two additional set of simulations. The first set of simulation uses intraspecific competition coefficients drawn from a normal distribution with mean 0.10 and standard deviation 0.01. While most resulting values would still ensure a frequency-dependent priority effect by positive frequency dependence (all intraspecific competition coefficients smaller than the baseline interspecific competition coefficient, ($\frac{B}{2} = 0.1125$), we expect that the general strength of frequency-dependent priority effects to be weaker. The second set of simulation uses a mean intraspecific competition coefficient of 0.15, much higher than baseline interspecific competition coefficient (0.1125). Without trait-dependent priority effects, this choice will lead to coexistence. For simplicity, we only report the most diagnostic results of these simulations, population dynamics within a patch and dispersal-diversity relationships.

Overall, both population dynamics and dispersal-diversity relationships change remarkably under different values of intraspecific competition without trait-dependent priority effects. With trait-dependent priority effects, these patterns do not change with intraspecific competition and initial scenarios. This is because with trait-dependent priority effects, outcomes of competition are mostly driven by emergence times.

#### Medium Intraspecific Competition

Population dynamics under Scenario 1 is similar to those observed with a lower intraspecific competition, but without trait-dependent priority effects the population trajectories are more clustered because of the weaker frequency-dependent priority effect (Figure S8). Under Scenario 2, and without trait-dependent priority effects, the initial resident (species 3) still remains mostly dominant in this patch at a low dispersal rate ($r = 0.01$), although in some simulations, a higher influx of other species can overcome the initial numeric advantage of species 3 (Figure S9). This is due to the weakened frequency-dependent priority effects of the resident species. This pattern is more obvious when dispersal rate is high ($r = 0.1$). In contrast, population dynamics with trait-dependent priority effects do not change much with higher intraspecific competition coefficients, as we still observe high turnovers of community composition within each patch.

Dispersal-diversity relationships with a higher intraspecific competition resembles those presented in the main text: dispersal strongly decreases beta and gamma diversity while creating a hump-shaped pattern for alpha diversity without trait-dependent priority effects; with trait-dependent priority effects, alpha and gamma diversity increases, and beta diversity remains constant with dispersal (Figure S10). Under Scenario 1 (Equal Initials) the hump-shaped dispersal-diversity relationships without trait-dependent priority effects are much weaker than those observed with the lowest intraspecific competition in the main text. This is because slightly higher intraspecific competition coefficients lower the presence of frequency-dependent priority effects, leading to more competitive exclusion and therefore lower diversity, especially under Scenario 1.

#### High Intraspecific Competition

As expected, intraspecific competition coefficients higher than $\frac{B}{2} = 0.1125$ promote coexistence without trait-dependent priority effects. Under both Scenario 1 and 2, all species randomly fluctuate but none are extinct; this observation is not affected by dispersal rates (Figure S11, S12). With trait-dependent priority effects, population dynamics are similar to simulations with low and medium intraspecific competition coefficients. Alpha and gamma diversity in metacommunities without trait-dependent priority effects remains high regardless of dispersal rates, reflecting local and regional coexistence observed from population dynamics (Figure S13). With trait-dependent priority effects, alpha and gamma diversity is lower, but increases with dispersal. Beta diversity shows the opposite trend: without trait-dependent priority effects, beta diversity is low and increases with dispersal, while beta diversity in metacommunities with trait-dependent priority effects remains relatively high, at a similar range observed with low and medium intraspecific competition (Figure S13). All above dispersal-diversity relationships are robust to initial conditions.

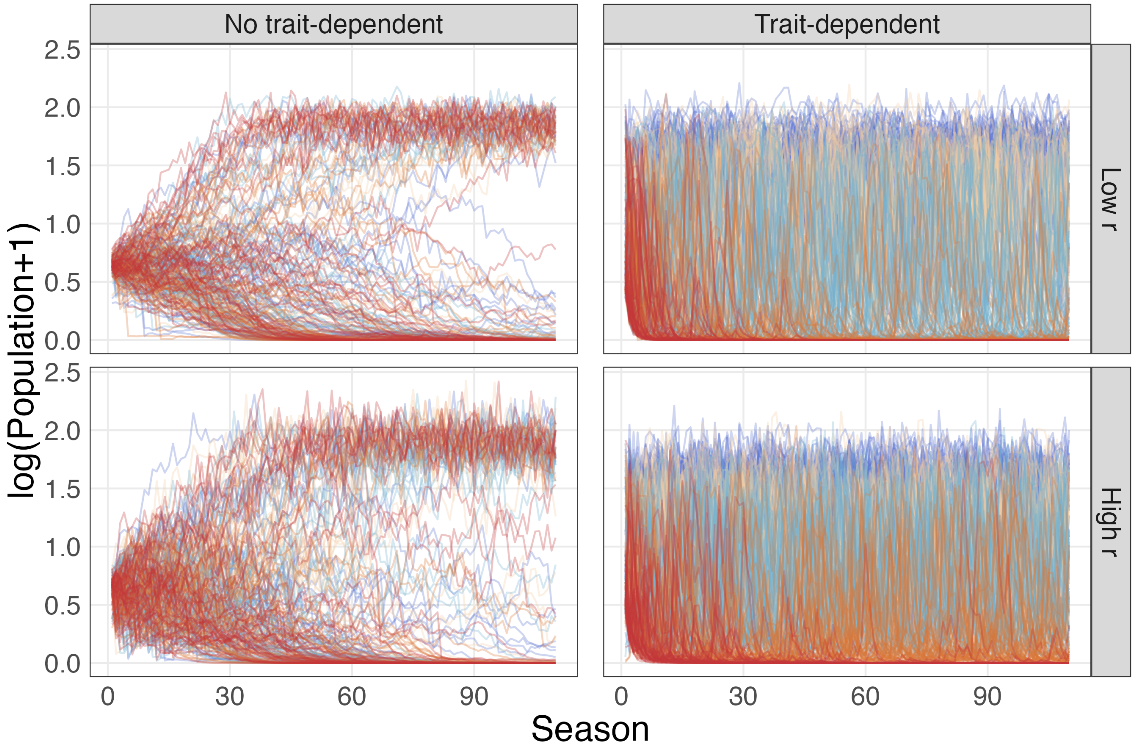


Figure S8. Example of population dynamics of one randomly selected patch under Scenario 1 (Equal Initials) and different dispersal rates, using an intermediate mean intraspecific competition, with and without trait-dependent priority effects. Line colors represent species. The left column shows dynamics without trait-dependent priority effects, and the right column shows dynamics with trait-dependent priority effects. Rows show results for low (0.01) and high (0.1) dispersal rates ($r$). Each line of the same color in a panel represents the population dynamics of that species from one out of 50 simulations. All population densities are transformed by natural log. See Table 1 for other parameters used.

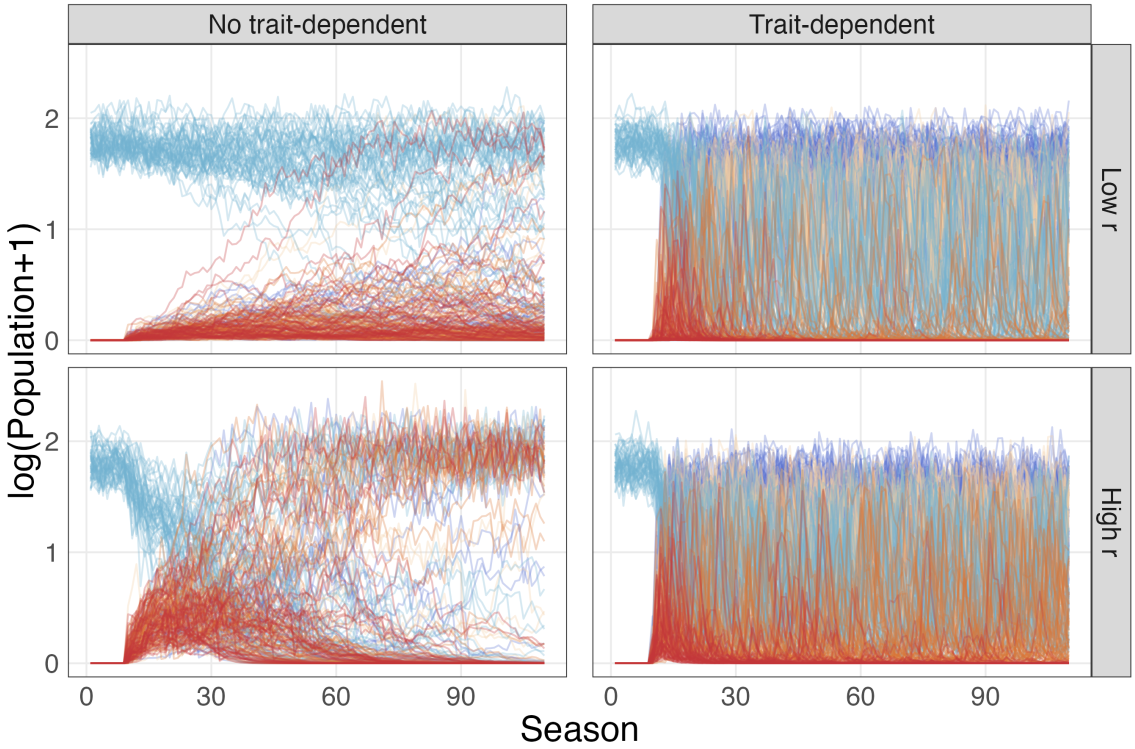


Figure S9. Example of population dynamics of one randomly selected patch under Scenario 2 (First Colonizer) and different dispersal rates, using an intermediate mean intraspecific competition, with and without trait-dependent priority effects. Line colors represent species. The left column shows dynamics without trait-dependent priority effects, and the right column shows dynamics with trait-dependent priority effects. Rows show results for low (0.01) and high (0.1) dispersal rates ($r$). Each line of the same color in a panel represents the population dynamics of that species from one out of 50 simulations. All population densities are transformed by natural log. See Table 1 for other parameters used.

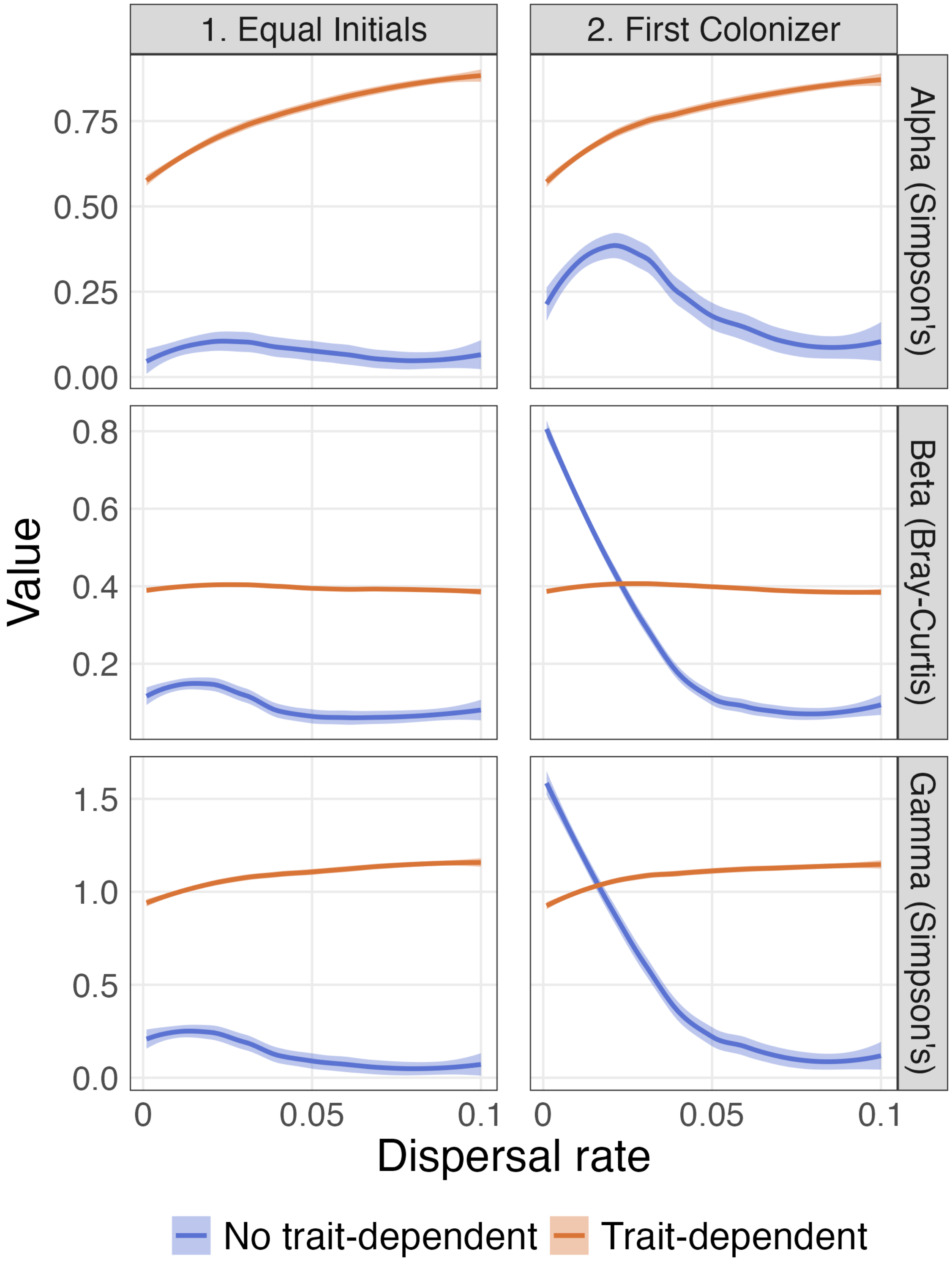


Figure S10. Relationship of alpha, beta, and gamma diversity and dispersal rates under the two initial scenarios and an intermediate mean intraspecific competition, with or without trait-dependent priority effects. Left and right columns show dispersal-diversity relationships under Scenario 1 (Equal Initials) and 2 (First Colonizer), respectively. Each line is rendered from data of all 50 simulations. See Table 1 for other parameters used.

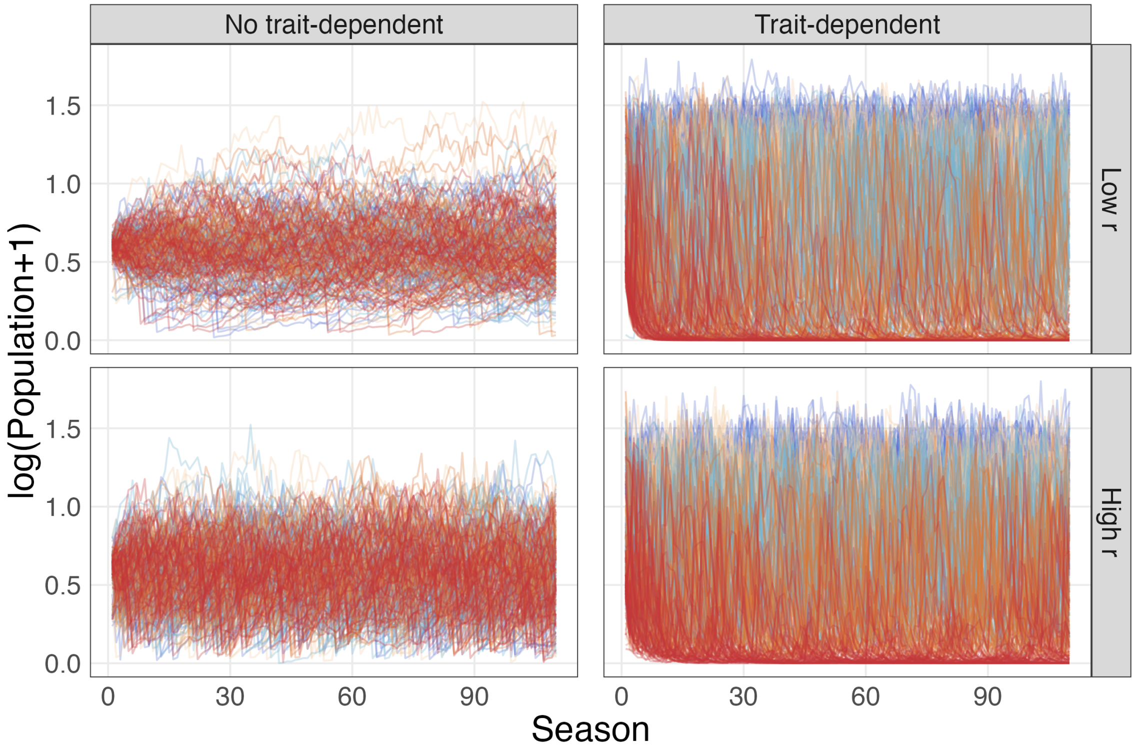


Figure S11. Example of population dynamics of one randomly selected patch under Scenario 1 (Equal Initials) and different dispersal rates, using a high mean intraspecific competition, with and without trait-dependent priority effects. Line colors represent species. The left column shows dynamics without trait-dependent priority effects, and the right column shows dynamics with trait-dependent priority effects. Rows show results for low (0.01) and high (0.1) dispersal rates ($r$). Each line of the same color in a panel represents the population dynamics of that species from one out of 50 simulations. All population densities are transformed by natural log. See Table 1 for other parameters used.

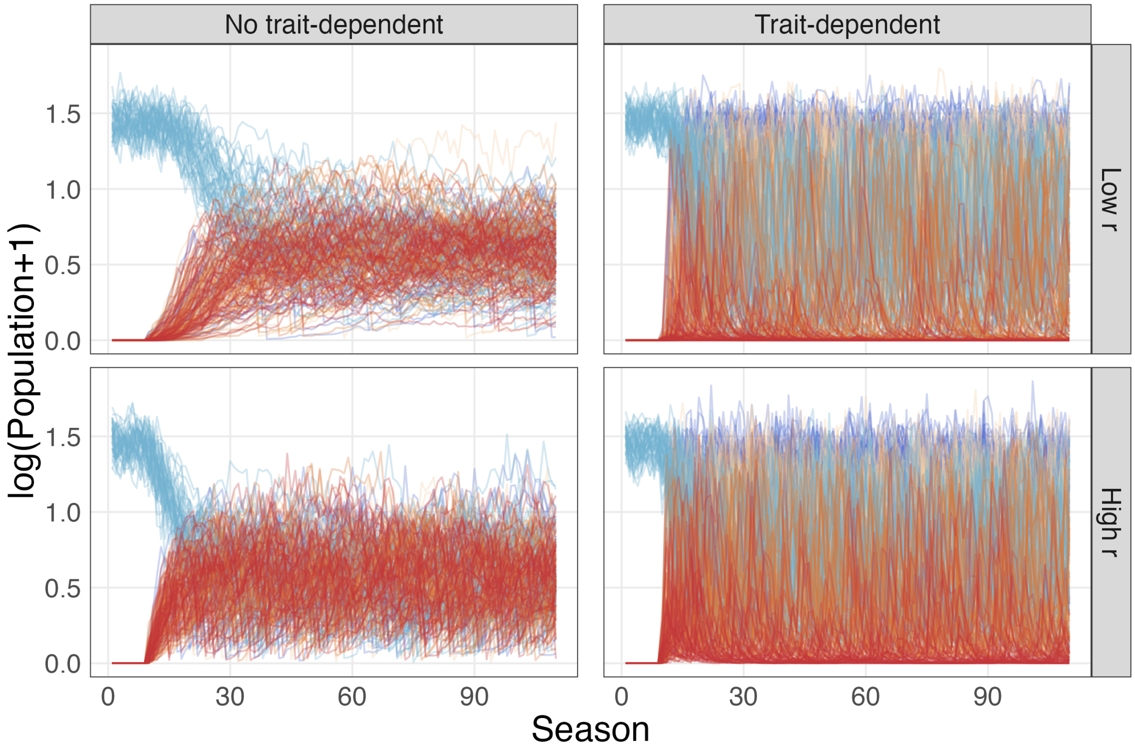


Figure S12. Example of population dynamics of one randomly selected patch under Scenario 2 (First Colonizer) and different dispersal rates, using a high mean intraspecific competition, with and without trait-dependent priority effects. Line colors represent species. The left column shows dynamics without trait-dependent priority effects, and the right column shows dynamics with trait-dependent priority effects. Rows show results for low (0.01) and high (0.1) dispersal rates ($r$). Each line of the same color in a panel represents the population dynamics of that species from one out of 50 simulations. All population densities are transformed by natural log. See Table 1 for other parameters used.

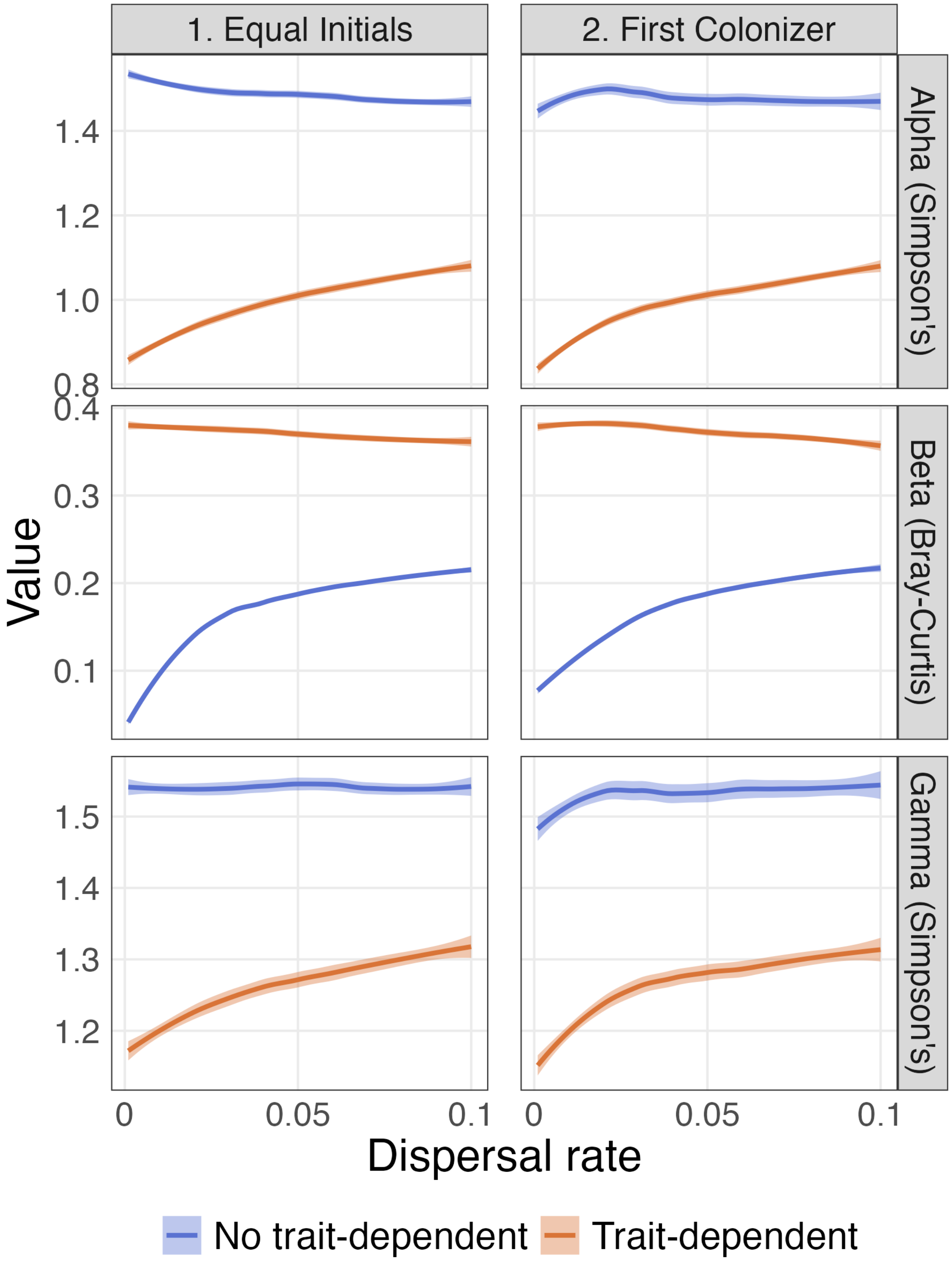


Figure S13. Relationship of alpha, beta, and gamma diversity and dispersal rates under the two initial scenarios and a high mean intraspecific competition, with or without trait-dependent priority effects. Left and right columns show dispersal-diversity relationships under Scenario 1 (Equal Initials) and 2 (First Colonizer), respectively. Each line is rendered from data of all 50 simulations. See Table 1 for other parameters used.
